# Dynamical System Modeling to Simulate Donor T Cell Response to Whole Exome Sequencing-Derived Recipient Peptides: Understanding Randomness In Clinical Outcomes Following Stem Cell Transplantation

**DOI:** 10.1101/088831

**Authors:** V Koparde, B Abdul Razzaq, T Suntum, R Sabo, A Scalora, M Serrano, M Jameson-Lee, C Hall, D Kobulnicky, N Sheth, J Sampson, J Reed, C Roberts, R Qayyum, G Buck, M Neale, A. Toor

## Abstract

The quantitative relationship between the magnitude of variation in minor histocompatibility antigens (mHA) and graft versus host disease (GVHD) pathophysiology in stem cell transplant (SCT) donor-recipient pairs (DRP) is not established. In order to elucidate this relationship, whole exome sequencing (WES) was performed on 27 HLA matched related (MRD), & 50 unrelated donors (URD), to identify nonsynonymous single nucleotide polymorphisms (SNPs). An average 2,463 SNPs were identified in MRD, and 4,287 in URD DRP (p<0.01); resulting peptide antigens that may be presented on HLA class I molecules in each DRP were derived *in silico* (NetMHCpan ver2.0) and the tissue expression of proteins these were derived from determined (GTex). MRD DRP had an average 3,670 HLA-binding-alloreactive peptides, putative mHA (pmHA) with an IC50 of <500 nM, and URD, had 5,386 (p<0.01). To simulate an alloreactive donor cytotoxic T cell response, the array of pmHA in each patient was considered as an *operator* matrix modifying a hypothetical cytotoxic T cell clonal *vector* matrix; each responding T cell clone’s proliferation was determined by the logistic equation of growth, accounting for HLA binding affinity and tissue expression of each alloreactive peptide. The resulting *simulated* organ-specific alloreactive T cell clonal growth revealed marked variability, with the T cell count differences spanning orders of magnitude between different DRP. Despite an estimated, uniform set of constants used in the model for all DRP, and a heterogeneously treated group of patients higher total and organ-specific T cell counts were associated with cumulative incidence of GVHD in recipients in Cox proportional hazard models. In conclusion, exome wide sequence differences and the variable alloreactive peptide binding to HLA in each DRP yields a large range of possible alloreactive donor T cell responses. Our findings also help understand the apparent randomness observed in the development of alloimmune responses.

## Introduction

Over the last four decades there have been substantial strides made in improving the clinical outcomes following allogeneic stem cell transplantation (SCT). Nevertheless, poor outcomes such as relapse and graft versus host disease (GVHD) remain difficult to predict in individuals because of the variability observed in the incidence of alloreactivity following both HLA-matched and HLA-mismatched SCT [1, 2, 3, 4, 5, 6, 7, 8, 9]. Considering that disease responses in allogeneic SCT are often linked to the development of GVHD, it is important to understand the biological basis for the incidence of alloreactivity in unique SCT donor-recipient pairs (DRP) [10]. It is well known that relapse and GVHD occurrence are a function of the magnitude of donor-derived immune reconstitution; however, in contrast to the occurrence of alloreactivity in cohorts of SCT recipients, immune reconstitution in individual DRP shows many characteristics of a dynamical system, such as logistic growth kinetics and power law distribution of clonal frequencies [11, 12, 13, 14, 15]. This implies that if clinical outcomes can be modeled as a function of donor derived immune reconstitution in unique transplant DRP, it will become possible to modify the system to optimize clinical outcomes. When fully developed such a model may allow *a priori* simulation of SCT with different donors, potentially leading to personalized immunosuppressive therapy.

To develop a simple model to simulate different donor T cell responses to unique recipient antigens, the array of antigens presented in each DRP would have to be considered. In HLA Matched SCT, recipient *minor histocompatibility antigens* (mHA) are presented on the HLA molecules to donor T cells. The response of these donor T cells to recipient antigens may be modeled as a dynamical system, using quantitative rules that govern repertoire evolution. To accomplish this the antigenic background in a DRP may be similarly described mathematically as a component of this dynamical system. These antigenic differences in a given transplant DRP may be *partially* determined using whole exome sequencing (WES) of SCT donor and recipient DNA. In previous work WES has demonstrated that there is a large library of non-synonymous single nucleotide polymorphisms (nsSNP) present in the recipient but absent in the donor, from which an equally large array of recipient-nsSNP-derived-peptides may be determined. These peptides have different amino acid sequences in each DRP [16, 17]. These immunogenic mHA bound to the ‘matched’ HLA molecules, may trigger donor T cell activation and proliferation. In aggregate these polymorphisms constitute an *alloreactivity potential* between the specific donors and recipients. Prior studies using dynamical system modeling of the T cell response to this mHA array shows marked variation in the simulated T cell response, which suggests that the alloreactivity potential varies considerably in different DRP [18]. Similar observations reporting association of polymorphisms in the peptide regions of HLA class I molecules support the premise outlined above [19].

Nevertheless, alloreactivity is a complex clinical state, where patients may have variable manifestations of GVHD impacting different organ systems to varying extent. In this paper the impact of tissue-specific-expression of the proteins from which the putative mHA are derived in different individuals is examined to measure its variability between unique transplant DRP. This is done using a T cell vector-mHA operator system previously developed [18] with a uniform set of conditions employed to simulate a hypothetical CD8+ T cell response to the *in silico* derived mHA-HLA class I array in 77 HLA matched SCT DRP.

## Methods

### Patients

Whole exome sequencing (WES) was performed on previously cryopreserved DNA obtained from donors and recipients of allogeneic SCT. Permission for this retrospective study was obtained from the Virginia Commonwealth University’s institutional review board, and patients who underwent transplantation between 2010 and 2014 were retrospectively selected for this analysis. Patients had undergone either 8/8 (n=67) or 7/8 (n=10) HLA-A, B, C and DRB1 matched related (MRD; n=27) or unrelated (MUD; n=50) or haploidentical (n=1) SCT according to institutional standards at Virginia Commonwealth University (VCU) (Supplementary Table 1). HLA matching had been performed using high resolution typing for the unrelated donor SCT recipients; and intermediate resolution typing for class I, and high resolution typing for class II antigens for related donor recipients. A variety of different conditioning and GVHD prophylaxis regimens were used in the patients.

**Table 1.** Exome sequencing and peptide results by donor type (n=77)

|  | MRD<sup>a</sup><br>(n=27) | MUD<br>(n=50) | P-value<sup>b</sup> |
| --- | --- | --- | --- |
| **Exome sequencing SNP differences** |  |  |  |
| Synonymous | 2,716 ± 703 | 4,821 ± 1358 | <0.01 |
| Non-synonymous | 2,463 ± 603 | 4,287 ± 1154 | <0.01 |
| Conservative | 847 ± 218 | 1,476 ± 402 | <0.01 |
| Non-conservative | 1,616 ± 388 | 2,811 ± 754 | <0.01 |
| **HLA-Presented Peptides** |  |  |  |
| Presented Peptides (IC50 <500 nM) | 3,670 ± 1551 | 5,386 ± 2136 | <0.01 |
| Strong HLA Binders (IC50 <50 nM) | 852 ± 373 | 1,160 ± 575 | <0.01 |

<sup>a</sup> Excludes haploidentical patient
<sup>b</sup> Mean values ± SD. P values determined using t-test for Equality of Means

#### Whole Exome Sequencing

Nextera Rapid Capture Expanded Exome Kit was used to extract exomic regions from the deidentified DNA samples, which were then multiplexed and sequenced on an Illumina HiSeq 2500 to achieve an average coverage of ^∼^90X per sample. 2X100 bp sequencing reads were then aligned to the human reference genome using BWA aligner. Duplicate read alignments were detected and removed using Picard tools. Single nucleotide polymorphisms (SNPs) in both the donor and recipients’ exomes were determined using GATK HaplotypeCaller walker. GATK best practices were then implemented to filter and recalibrate the SNPs; and store them in variant call file (VCF) format. To identify SNPs unique to the recipient and absent in the donor the results from the GATK pipeline in VCF format were then parsed through the in-house TraCS (Transplant pair Comparison System) set of perl scripts. TraCS traverses through the genotypes of the called SNPs, systematically excluding identical SNPs or editing them to align with the graft-versus-host (GVH) direction thereby generating a new VCF with SNPs for a particular DRP in the GVH direction (SNP present in the recipient, absent in the donor; R^+^/D^-^).

The SNPs in this VCF are then annotated either as synonymous or non-synonymous using Annovar. The corresponding amino acid polymorphisms along with flanking regions of each protein are then extracted using Annovar to build peptide libraries of 17-mers for each DRP, with the SNP encoded AA occupying the central position. This library is further expanded by sliding a 9-mer window over each 17-mer such that the polymorphic amino-acid position changes in each 9-mer. The HLA class I binding affinity and IC50 values, which quantify the interactions between all these 9-mers for each DRP and all six HLA class I donor molecules (HLA-A, B and C), NetHMCpan version 2.8 was run iteratively in parallel mode on a linux cluster using custom python scripts. Parsing the NetMHCPan output, unique peptide-HLA combinations present in the recipient but not in the donor, i.e., possessing a GVHD vector, were identified and organized in order of declining mHA-HLA affinity. IC50 (nM) indicated the amount of peptide required to displace 50% of intended or standard peptides specific to a given HLA. Binding affinity is inversely related to IC50, such that, smaller the IC50 value, the stronger is the affinity. The variant alloreactive peptides with a cutoff value of IC50 ≤500 are included in the analyses presented here.

The Genotype-Tissue Expression (GTEx) portal V6 has publicly available expression level information (Reads/kilobase of transcripts/million mapped reads, RPKM values; http://www.gtexportal.org/home/) for a variety of human tissues over a large number of genes. Since the gene-ids for the proteins that generate the peptides in our DRP peptide library are known, the RPKM values from the GTEx portal for the specific gene across the whole array of tissues of interest can be parsed in, namely, skin, lung, salivary gland, esophagus, small intestine, stomach, colon and liver. In the DRP where a male recipient had been transplanted from a female donor (n=17), full length available sequences of all proteins encoded by the Y chromosome were curated from NCBI. These sequences were then computationally split into 9-mer peptides and their respective binding affinities and IC50 values for the relevant donor HLA antigens were determined *in silico* using the NetMHCpan software as described earlier. These peptides were then appended to the corresponding DRPs peptide library for all subsequent analyses.

#### Computational Methods: Dynamical System Modeling Of T Cell Response To Putative mHA

To simulate the cytotoxic T cell response to HLA class I bound antigens, it was postulated that the immune effectors and their antigenic targets constitute a *dynamical system*. This was modeled as an *iterating* physical system which evolves over time, such that the state of the system at any given time (*t*), depends on the preceding state of the systems. The system in this instance was comprised of all the known peptide HLA complexes and the hypothetical donor T cell response to the same. In this system, each T cell clone responding to its target antigen will proliferate, conforming to the *iterating logistic equation* of the general form

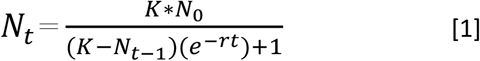

In this equation: *N*_*0*_, is the T cell count at the outset (for the calculations presented in this paper *N*_*0*_=1 at *t*=1); *N*_*t*_, is the T cell count at time *t* following transplant (*t* modeled as iterations); *N*_*t-1*_ represents the T cell count at the previous iteration; *K* is the T cell count at the asymptote (steady state conditions after infinite iterations), and represents the maximum T cell count the system would support (carrying capacity); *r* is the growth rate. The T cell population, at time *t, N*_*t*_, depends on the preceding T cell populations, such that

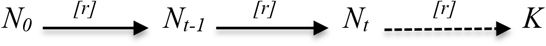

Expanded to the entire set of T cells responding to all recipient antigens, the final steady state population of all the T cell clones 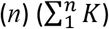, and its clonal repertoire may be studied for associations with clinical outcomes. In such a model, the parameter *r* can have either positive or negative values depending on whether the ambient cytokine milieu, either stimulates (-*r*) or suppresses (+*r*) growth. Simulating this logistic dynamical system consists of repeated calculations, where the result for each iteration gives the population N_t_, as a function of time and becomes the input variable for the next calculation N_*t*+1_. This system behaves in a non-linear fashion, demonstrating sigmoid population growth constrained by feedback [18, 20, 21].

#### Matrices To Model The T Cell Clonal Repertoire: Vector Spaces And Operators

A system of matrices is utilized to apply the logistic equation to describe the evolution of all the unique T cell clones present in an individual at a given time, in other words, to *simulate* the evolution of the entire T cell clonal repertoire responding to the WES derived mHA-HLA library for a transplant DRP. For this simulation, T cell clones present in an individual are considered as a set of individual *vectors* in the immune *phase space* of that individual. The descriptor, *vector*, in this instance is used to describe the entire set of T cells, with the individual T cell clones representing components of this vector [22]. In this vector, T cell clonality represents direction (since it determines antigen specificity) and T cell clonal frequency, the magnitude of individual vector components. The sum of the elements of these vectors will represent the overall T cell vector and its direction; in this simulation, recipient-directed alloreactivity. The T cell vectors may take many different configurations in an individual, and the entire range of possible vector configurations constitutes the *immune phase space* for that individual. The T cell vector may be represented mathematically as a single row matrix, with T cell clonal frequencies *TC_1_, TC_2_, TC_3_…. TC_n_*. This T cell clonal matrix is represented by the term, 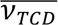. The frequency of each TC is a natural number, and the direction, its reactivity, represented by the clonality. Phase space will then describe all the potential T cell clonal frequency combinations possible, within a specific antigenic background, with all the potential directions of the T cell reactivity. In the case reported here, the vectors are all in the direction of graft vs. host alloreactivity, since they proliferate after encountering mHA-HLA. In contrast, the infused T cell clonal repertoire will represent a steady state T cell clonal frequency distribution in the normal donor, presumably made up of mostly self-tolerant, pathogen specific T cells, and without any auto-reactive T cells. For the computations reported here, each T cell clone has a *N*_*0*_ value of 1, and is alloreactive.

The T cells infused with the transplant encounter a new set of antigens (both recipient and pathogen). These antigens presented by the recipient (or donor-derived) antigen presenting cells constitute an *operator*, a matrix of targets in response to which the donor T cells proliferate [23]. This operator changes the magnitude and direction (clonal dominance) of the infused donor T cell repertoire, as individual T cell clones in the donor product grow or shrink in the new HLA-antigen milieu, transforming the T cell vector. The putative mHA making up the alloreactivity potential constitute a matrix, termed an *alloreactivity potential operator*, 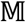_APO_. This operator incorporates the binding affinity of the variant peptide-HLA complexes encountered by donor T cells in the recipient. Following SCT, the donor T cell clonal frequency changes depending on the specificity of the TCR and the abundance and reactivity of the corresponding antigen. This corresponds to the operator modifying the <u>original</u> *donor* T cell vector, 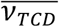 as the system goes through successive iterations to a *recipient* T cell vector, 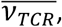 according to the following relationship

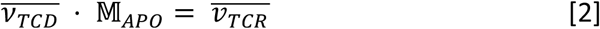

#### Applying the Logistic Growth Equation to Vector-Operator Systems

A central assumption in this model is that the steady state TC clonal frequency (*K*) of specific TCR bearing clones, and their growth rates (*r*), are proportional to the binding affinity of their target mHA-HLA complexes. This is because the strength of the binding affinity will increase the likelihood of T cell-APC interactions occurring, thus influencing the driving rate of this interaction. This is *approximately* represented by the reciprocal of the *IC50* for that specific complex. While each TCR may recognize another mHA-HLA complex with equal or lesser affinity, for the sake of simplicity, an assumption of non-recognition of other mHA-HLA complexes was made for this model. This makes it possible to specify an *identity matrix*, Matrix A, with unique mHA-HLA complexes for a SCT-DRP and the corresponding TCR, where *TCR*_*x*_ binds *mHA*_*x*_*-HLA (relative* binding affinity, 1) and not others (*relative* binding affinity, 0). In addition to the assumption that there is a first signal of TCR recognition, a second signal for immune responsiveness is assumed in this matrix.

<u>Matrix I.</u> Effect of alloreactivity operator 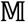_APO_ on T cell vector 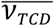. Successive iterations (*t, t+1 etc*.) modify the vector to 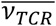. In this simplified model, *mHA*_*x*_*-HLA* only binds *TC*_*x*_ and so on. Matrix below represents a single iteration *t*.

| SCT | $M_{APO}$ | | | | |
| --- | --- | --- | --- | --- | --- |
|  |  | mHA<sub>1</sub> HLA | mHA<sub>2</sub> HLA | mHA<sub>3</sub> HLA | mHA<sub>n</sub> HLA |
| $\overline{v_{TCD}}$ | $TC_1$ | 1 | 0 | 0 | 0 |
| $\downarrow t$ | $TC_2$ | 0 | 1 | 0 | 0 |
| $\overline{v_{TCR}}$ | $TC_3$ | 0 | 0 | 1 | 0 |
| | $TC_n$ | 0 | 0 | 0 | 1 |

The *alloreactivity matrix* modifies the donor T cell clonal vector infused with an allotransplant, *mapping* it to the recipient T cell vector, as the T cell clones with unique TCR encounter the corresponding mHA-HLA complexes they proliferate conforming to the logistic equation [Equation 1]. In the logistic equation *K* for each T cell clone will be proportional to the approximate binding affinity (*af* = 1/*IC*50) of the corresponding mHA-HLA complex (*K^af^*). In this model, the parameter *r* is also a function of the binding affinity, and reflects the effect of the cytokines driving T cell proliferation. Equation 1 must therefore be modified for unique TC clone TC*x*, responding to *mHAx-HLA*, as follows:

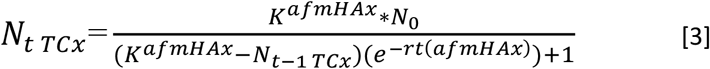

In this equation the parameter *afmHAx* represents the binding affinity of the mHA-HLA complex, *x*, given by the expression 1/*IC50x*. This equation gives instantaneous T cell counts in response to antigens presented when *N*_*0*_ equals 1, regardless of tissue distribution. In the alloreactivity, vector-operator identity matrix, the values 1 or 0 in each cell are multiplied by the product of Equation 3 for each T cell clone. This is depicted in Matrix II.

<u>Matrix II.</u> Matrix illustrating a single iteration of the alloreactivity operator 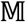_*APO*_ on T cell vector 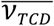. Each cell in the matrix calculates the value of *TC*_*x*_ in response to *mHA*_*x*_*-HLA*, final repertoire size (or magnitude of T cell response to antigen array) is determined by solving the matrix.

| SCT | $\mathbb{M}_{APO}$ | | | | |
| --- | --- | --- | --- | --- | --- |
|  |  | mHA<sub>1</sub> HLA | mHA<sub>2</sub> HLA | mHA<sub>3</sub> HLA | mHA<sub>x</sub> HLA |
| $\overline{V_{TCD}}$<br>$\downarrow t$<br>$\overline{V_{TCR}}$ | $TC_1$ | $N_{tTC1} = \frac{K^{afmHA1}}{(K^{afmHA1} - N_{t-1TC1})(e^{-rt(afmHA1)}) + 1}$ | 0 | 0 | 0 |
| | $TC_2$ | 0 | $N_tTC_2$ | 0 | 0 |
| | $TC_3$ | 0 | 0 | $N_tTC_3$ | 0 |
| | $TC_x$ | 0 | 0 | 0 | $N_tTC_x$ |

The T cell response to each mHA-HLA complex is determined over time, *t*, by iterating the system of matrix-equations. In this *alloreactivity matrix*, the IC50 of all the alloreactive peptides with a GVH direction (present in recipient, but absent in donor) constitutes the operator; the sum of *n* T cell clones 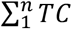, at each time point will represent the magnitude of the vector 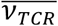 at that time *t*. In this system, when considering the effect on infused donor T cell vector, depending on antigen affinity, T cell clones present in abundance may be down-regulated if antigen is not encountered. Conversely, clones present at a low frequency may expand upon encountering antigen, *transforming* the vector over time from the original infused T cell vector. In summary, the alloreactivity operator determines the change in donor T cell vectors, following transplantation, in an iterative fashion *transforming* it to a new configuration, over time t, based on the mHA-HLA complexes encountered in the recipient and their affinity distribution (*and antigen abundance, vide infra)*. Therefore post-SCT immune reconstitution may be modeled as a process in which T cell clonal frequency vectors are iteratively multiplied by the minor histocompatibility antigen matrix operator and this results in transformation of the vector over time to either a GVHD-prone alloreactive or to a tolerant, pathogen-directed vector. This may be visualized as thousands of T cell clones interacting with antigen presenting cells, an example of an interacting dynamical system [11].

#### Competition between T Cell Clonal Populations

Each of the T cell clones behaves like a unique population, therefore *competition* with other T cell clones in the set of all T cell clones must be accounted for in the logistic equation to determine the magnitude of the unique clonal frequencies as the model iterates simulating T cell clonal growth over time. This may be done using the *Lotka-Volterra* model for *competing populations*, which accounts for the impact of population growth of multiple coexisting populations [18, 24, 25]. In the model presented here, this is accomplished by modifying the expression *N*_*t*-1_ in equation 3 for each clone, by taking the sum of *N*_*t*-1_ for all the other competing T cell populations when calculating *N*_*t*_ for each clone. Each clone’s *N*_*t*-1_ is weighted by a correction factor α, which is proportional to its interaction with the T cell clone being examined. Given the central role for the target mHA-HLA complex’s *IC50* in determining the T cell frequency for each clone, α for each clone is calculated by dividing the *IC50* of the competing T cell clone with the test clone (note the use of *IC50* instead of *1/IC50)*. This implies that T cell clones recognizing mHA-HLA complexes with a higher binding affinity will have a disproportionately higher impact on the growth of T cell clones binding less avid mHA-HLA complexes and vice versa. The resulting Matrix III, will have 1 on the diagonal, and values <1 above the diagonal, and >1 below it.

<u>Matrix III.</u> Matrix illustrating the relative effect of antigen binding affinity on the T cell clonal interaction between different clones. Successive cells in each row of the matrix calculate the effect of *TC_i_* on T cell being studied, *TC_x_*. This generates a weighting factor, α, which modulates the impact of population of *TC_i_* on the growth of *TC_x_*.

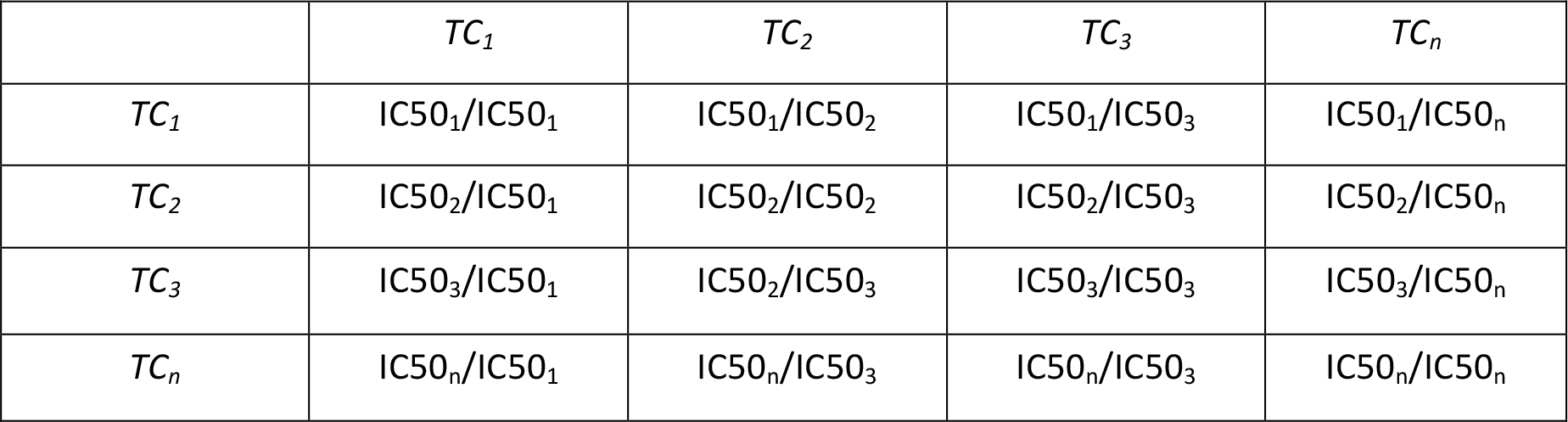

To account for *n* competing T cell populations, equation 3 is modified as follows

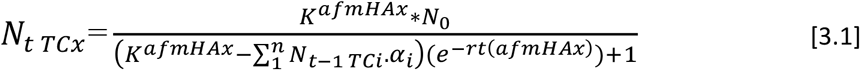

Where for the T cell clone *x* (*TC_x_*), *N_t_* depends on the sum of the *n* T cell clonal frequencies at the previous iteration. The α for each competing T cell clone *i*, with respect to TCx, modulates the effect of the magnitude of the *i*th T cell clone on *TC_x_*.

#### Accounting for tissue expression of proteins

The peptides discussed in the aforementioned derivation are generated from proteins expressed in target tissues, as such the level of protein expression will determine the magnitude of the peptide specific T cell response. The higher the protein expression, the more peptide molecules and the greater the HLA presentation with an ability to stimulate a larger T cell response (clonal frequency). Therefore, the parameter *K* in the above equations may be modeled as a multiple of the level of protein expression. The tissue expression of proteins (*Pexp)* is estimated by RNA sequencing techniques, and is measured in Reads Per Kilobase of transcript per Million mapped reads (RPKM). The values available from the public data base GTEx, range from 0 to >10^4^ and constitute a coefficient in the definition of K. This Modifies Equation 3.1,

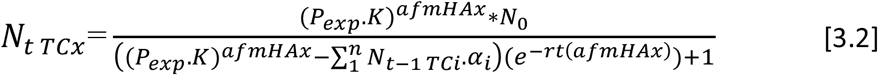
In equation 3.2, the tissue expression of the protein from which the target peptide, *mHAx* is derived is incorporated as a K multiplier when calculating a tissue-specific T cell response. Thus tissue-specific alloreactivity potentials may be simulated for each of the relevant GVHD target tissues, by substituting equation 3.2 into Matrix II. Solving the matrix yields an organ specific T cell response, given by the total T cell count, 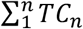.

In applying this model to exome sequence derived, alloreactive-peptide-HLA binding patient data an IC50 cutoff value of ≤500 nM, and an RPKM value of ≥1 were chosen to study the differences between patients. The T cell repertoire simulations were then performed in MATLAB (Mathworks Inc., Natick, MA), utilizing the above model (See Supplementary Mathematical Methods and Program).

The sum of all T cell clones 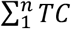 *TC* for each specific organ of interest and the grand total of the organs studied (termed sum of all clones) were used to represent the T cell vector magnitude of the alloreactive T cells following 500 iterations of the system for each patient.

#### Statistical Methods

Estimated T cell counts are summarized per organ and in total, with means and standard deviations, both overall and by GVHD classification. Acute GVHD was graded according to Glucksberg criteria and chronic GVHD by NIH consensus criteria. GVHD developing after DLI was not considered. The estimated T cell counts are compared between GVHD and Donor Type groups using the Wilcoxon rank sum test.

Cox proportional hazards models are used to examine the association between the estimated T cell counts and patient outcomes (GVHD, survival, relapse and relapse-free survival), with and without adjustment for age and patient gender and using robust standard error estimates. Cox proportional hazards analyses were performed in Stata 14 (StataCorp LP. College Station, TX). The MEANS, NPAR1WAY and PHREG procedures in the SAS statistical software program (version 9.4, Cary, NC, USA) are used for summaries and analyses. All estimated T cell counts are divided by 1000.

## Results

### Exome sequencing

Following de-identification, cryopreserved DNA samples from 78 donors and recipients of allogeneic SCT were sequenced, demographic details of the patients are given in Supplementary Table 1. Whole exome sequencing data analyzed using the TraCS program revealed a large number of nsSNPs with a GVH direction in each DRP (R^+^/D^-^). Recipients of MUD had a significantly larger number of SNPs when compared with MRD recipients with most SNPs being non-conservative (Table 1). In patients who either, did <u>or</u> did not develop GVHD, an average of 2,456 and 2,474 nsSNPs were identified in MRD recipients, while the corresponding numbers for MUD recipients were 4,536 and 3,845 nsSNPs (P=not significant (N.S.) for both). In the single haploidentical transplant recipient, there were 3,231 nsSNPs. Notably, the transition/transversion ratio in the WES data was as expected between 2.1 and 2.5 for the entire data set (Supplementary Fig 1).

### In silico derivation of mHA-HLA complexes from exome variation

The HLA class I binding peptides incorporating the amino acid coded by the nsSNP_GVH_ in each DRP were interrogated using NetMHCPan 2.8. These analyses yielded a large array of unique peptide-HLA complexes, putative or *potential* minor histocompatibility antigens (*pmHA)* for each pair. Utilizing all the nsSNP_GVH_, the MRD DRP had an average of 44,214 pmHA and the MUD DRP 77, 025 (P <0.01). A significantly larger number of HLA binding peptides with intermediate or high affinity were observed in MUD recipients when compared with MRD (Table 1). A similar number of peptides with an IC50 <500nM, 3,826 and 3,406 mHA, were recorded in MRD patients either with or without GVHD respectively; corresponding numbers of *pmHA* were 895 and 778 mHA with an IC50 <50nM for those patient groups (P=N.S.). For the MUD SCT pairs (n=50), there were 5,672 and 4,876 *pmHA* with an IC50 <500 nM, and 1,210 and 1,071 *pmHA* with an IC50 <50 nM in patients with or without GVHD (P=N.S.). Notably these data do not include the Y chromosome derived peptides presented by the class 1 HLA molecules in male recipients of female donors. The haploidentical DRP had 2,448 peptides with an IC50 of <500 nM out of a total 57,957 peptides; of these 484 had an IC50 <50 nM, when using the recipient HLA for performing the calculation.

### Simulating recipient tissue specific donor T cell responses

Following determination of the *pmHA* with various binding affinity distributions, the T cell responses to these antigens were simulated using the derived binding affinities in the T cell vector- alloreactivity operator model [Equations 2 & 3.2]. These simulations incorporated the tissue expression levels (in RPKM) from the GTEX portal for the proteins that generated the *pmHA*. Y-chromosome coded HLA binding peptides were also included in the DRP where a female donor had been used for a male recipient. A uniform set of values was programmed into the MATLAB software for other variables in equation 3.2, specifically, for each *pmHA* there was a single alloreactive T cell at the first iteration, i.e. *N*_*0*_= 1 at *t* = 1; *K* was set at 1,000,000 cells (this value substitutes for organ mass, lympho-vascular supply etc. without accounting for differences between organs), and *r* was set at 1.5 for all the simulations (in *vivo* this may vary with state of inflammation). All the *pmHA* were included in the simulations with competition. The program simulated donor T cell responses to skin, salivary gland, esophagus, stomach, small intestine, colon, liver, lung, blood, and finally T cell frequency for all these organs was summed.

Individual simulated T cell clonal growth was non-linear & variable depending on the HLA binding affinity of the peptide, as well as the tissue expression of protein (Figure 2A). As expected high levels of protein expression can compensate for low antigen binding affinity of the derived peptide, and vice versa. T cell clones responding to hi expression/hi binding affinity peptides, rose rapidly and dominated the emerging repertoire, while low expression/low binding affinity were initially suppressed, but recovered over time (iterations of the system). Accordingly, the overall clonal repertoire (number of clones, and cell number for each clone) in each DRP grew over time, with variable and unique growth kinetics (Figure 2B). Clonal repertoire responding to each organ conformed to power law distribution when sampled in four patients (patient number 2, 3, 4 & 5) (Supplementary figure 2). Individual T cell clonal growth for each organ’s alloreactive peptides was summed to give the total *simulated*, alloreactive T cell count for each organ in each DRP. When the number of clones constituting the overall and organ-specific T cell populations was determined following 500 iterations of the model, a high degree of variability was identified between both MRD and MUD DRP (Figure 2C).

**Figure 1.**
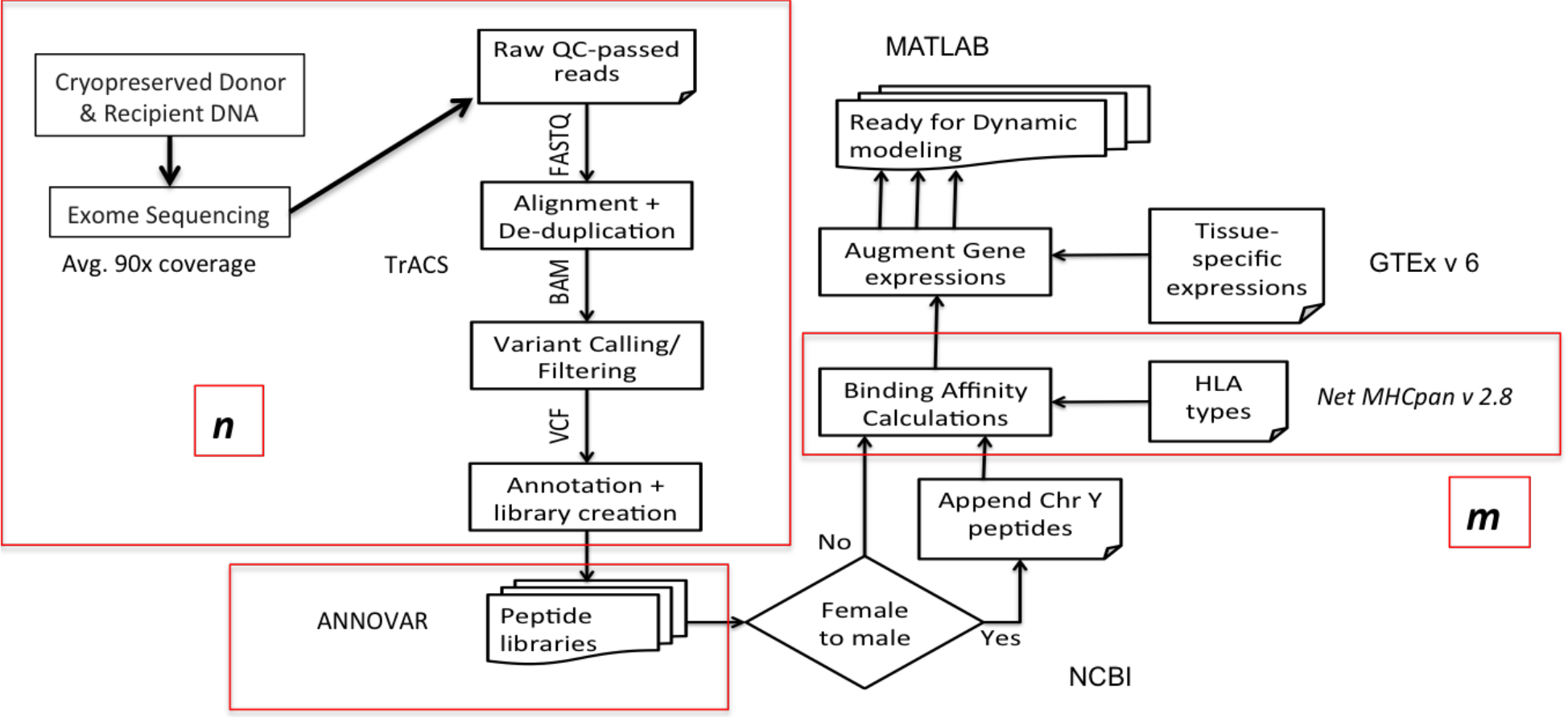
TRaCS, computational algorithm to determine tissue specific alloreactive putative peptide minor histocompatibility antigens (pmHA). Whole exome sequencing of cryopreserved donor and recipient DNA was performed, with an average coverage of 90x. The variable ***n*** refers to the number of nsSNP_GVH_ and ***m*** to pmHA. The variable ***m*** along with protein tissue expression level was then analyzed in MATLAB to determine T cell responses.

**Figure 2.**
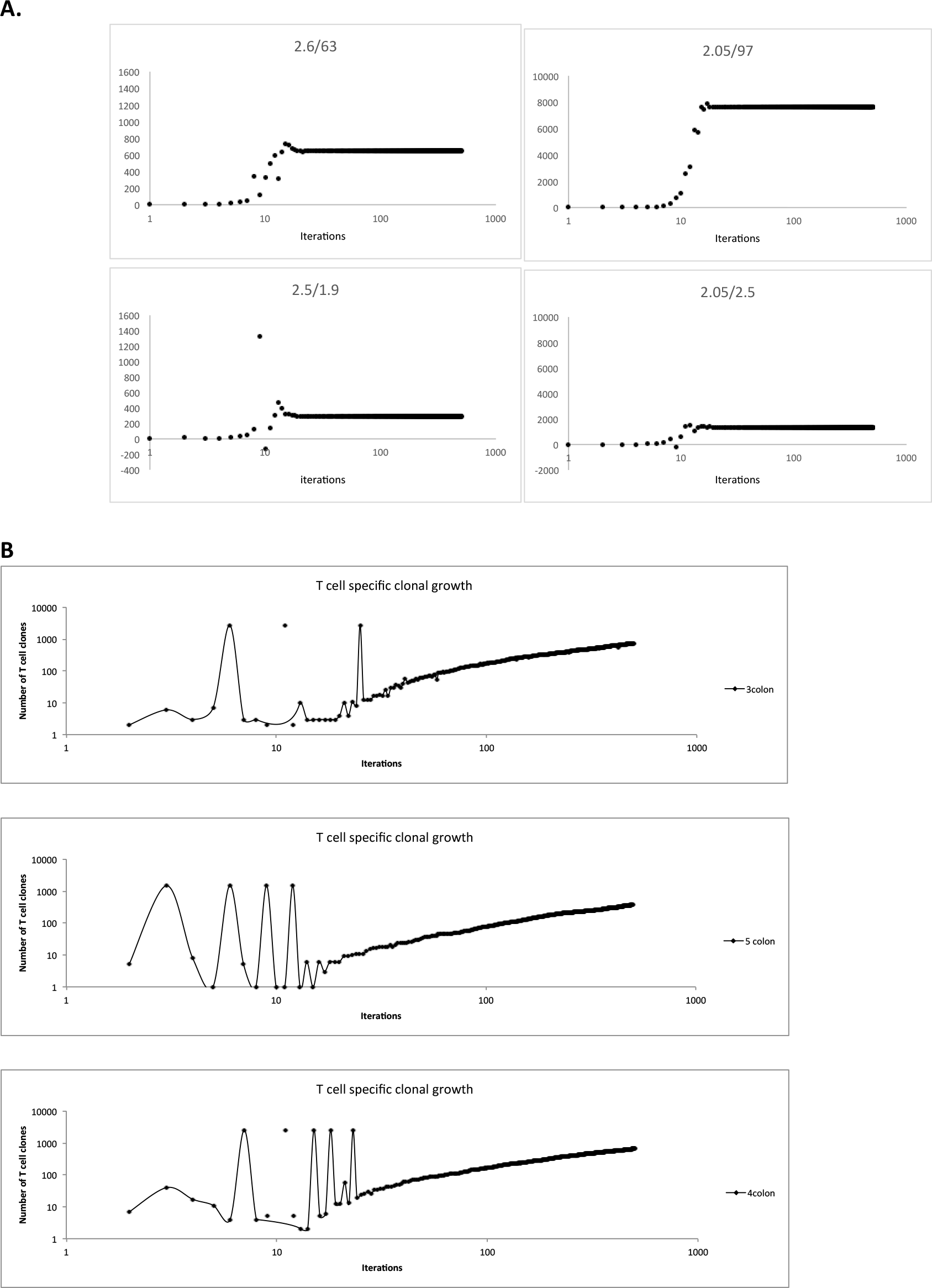
T cell clonal growth in the simulations. A. Individual T cell clone growth simulations accounting for peptide-HLA complex binding affinity and protein of origin tissue expression (IC50/RPKM). Increased T cell frequency (Y-axis) seen if the protein is expressed at a higher level. Different *pmHA* from a single patient/organ. B. Variable growth pattern of the number of clones in the simulations, number of clones rising over ‘time’ (iterations); T cell clonal growth in response to colonic alloreactive peptides depicted here.

**Figure 2C.**
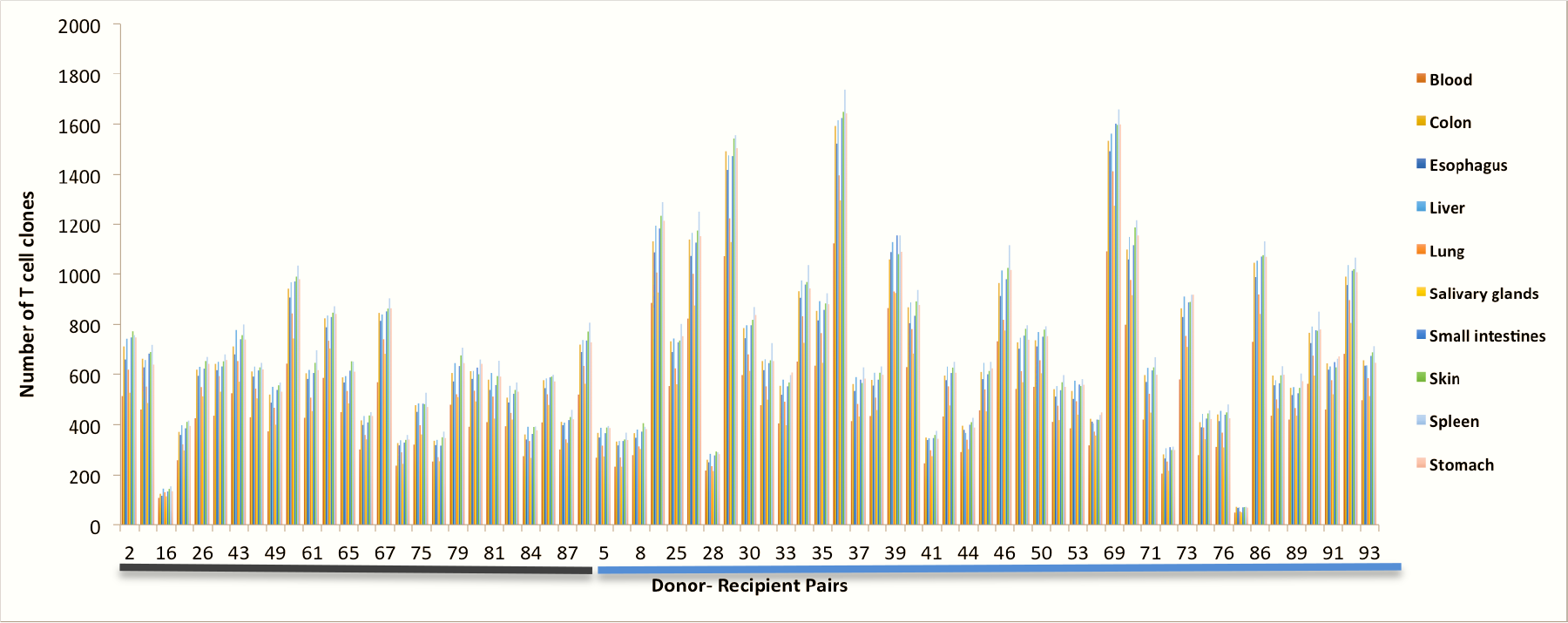
Number of T cell clones after 500 iterations, reflecting the number of high affinity peptides expressed in the tissues studied (GTEX). A non-significant trend towards a larger number of clones in MUD recipients is observed in this graph.

The average T cell count for each organ after 500 iterations (average of the iteration # 401-500) was determined (Figure 3A), and depicts the magnitude of organ-specific alloreactive T cell response that may be seen in different DRP, accounting for their HLA types and exome sequence variation. The differences observed in the organ-specific and overall simulated T cell responses to the HLA bound peptides between unique DRP spanned orders of magnitude. The simulated organ-specific T cell counts responding to the alloreactivity operator demonstrate this variability in both HLA matched related and unrelated donors, both between different DRP (Figure 3B) and to a lesser extent within each DRP (Figure 3C). This is true for individual organ simulations, as well as the sum of these simulations (Figure 3D), which reflects the variation in the organ specific T cell responses.

**Figure 3A.**
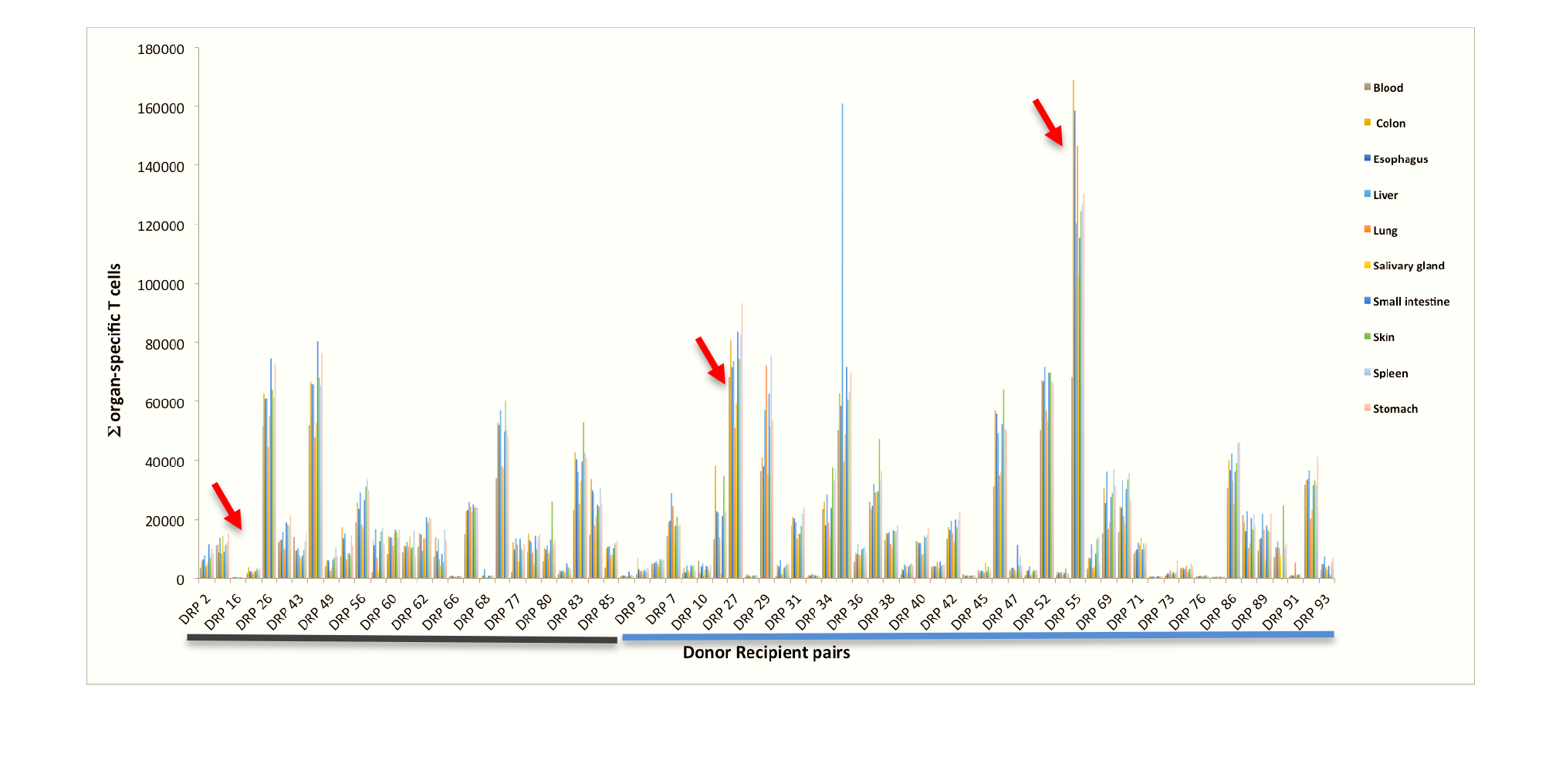
Tissue specific simulated alloreactive T cell counts after 500 iterations in recipients of MRD (Black line) and unrelated donors (Blue line). These values represent the sum of the entire T cell vector responding to the alloreactivity operator matrix of mHA-HLA complexes for a specific organ 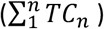. Arrows denote individuals with T cell responses differing by orders of magnitude. Values were obtained by calculating the average for iteration number 401-500 due to variability from competition.

**Figure 3B & 3C.**
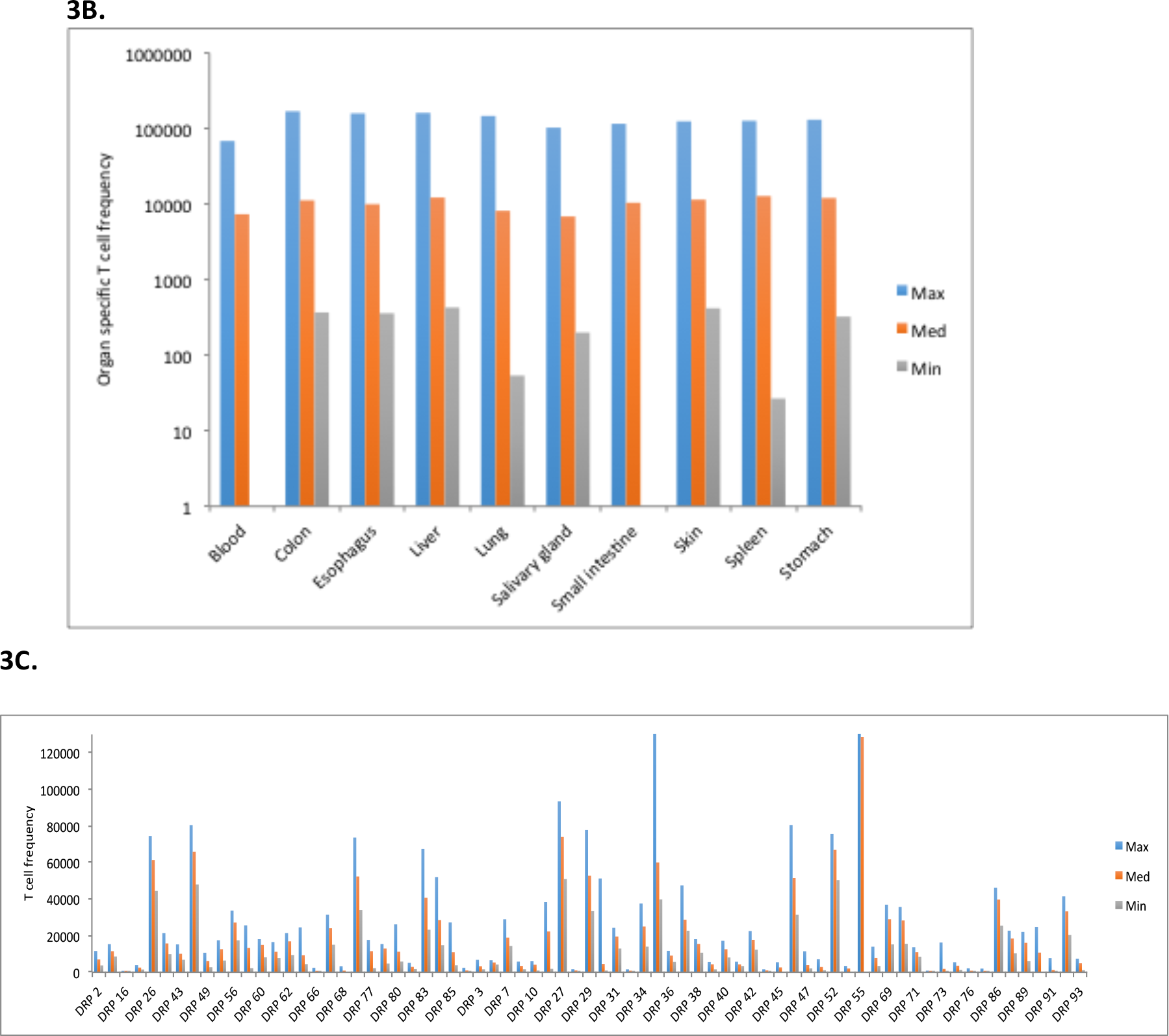
Variation in the simulated alloreactive T cell counts observed for each organ examined between patients (3B) and within patients (3C). Note Log scale used in Figure 3B, but not in 3C (Y axis truncated at 130000).

**Figure 3D.**
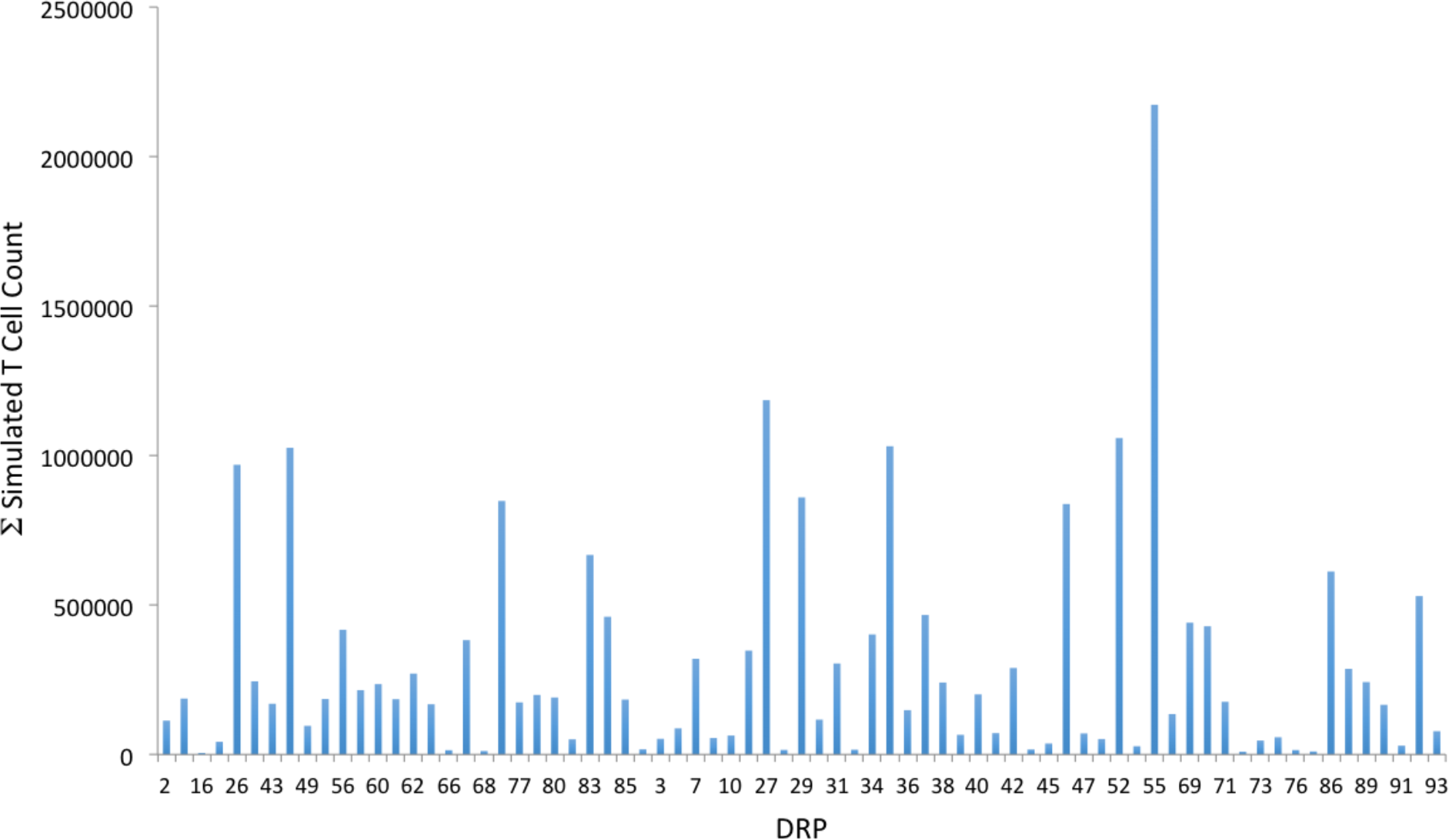
Total, simulated alloreactive T cell counts in recipients of MRD and unrelated donors after 500 iterations (average for iteration number 401-500 due to variability from competition). P value for magnitude difference, >0.3

When the sum of all organ specific T cell clones (Σ TC) was examined in different donor types, no significant differences were observed, however, while the differences were not statistically significant between groups, there was a trend supporting quantitative differences between different donor types. Simulated T cell counts are summarized by donor type in Supplementary Table 2, where for every organ, and overall, the mean T cell counts and estimated variabilities are larger in patients with unrelated donors than they are in patients with matched related donors, however these differences were not significant in this cohort (all p-values > 0.3). No statistically significant differences were observed when biological features such as gender mismatch (F to M: 270,613 (n=17), M to F: 135,064 (n=14) vs. gender match (F to F: 180,574 (n=16), M to M: 114,932 (n=26), and racial difference in the DRP (AA to AA: 113,245 (n=13), vs. C to C: 215,337 (n=47), vs. race mm: 57,938 (n=6)) were studied. This corroborates well with the relatively weak effect of these biological differences in the development of GVHD in the setting of HLA matching. [26, 27]

**Table 2.** Cox Proportional Hazards Model for GVHD association with simulated organ specific T cell counts, with and without adjustment for recipient age and gender, using robust S.E. estimates.

ORGAN
|  | Un-adjusted HR (95% CI) | P-value | Adjusted HR (95% CI) | P-value |
| --- | --- | --- | --- | --- |
| **Saliva** | 1.012 (1.000 to 1.027) | 0.070 | 1.013 (1.001 to 1.026) | **0.028** |
| **Colon** | 1.009 (1.000 to 1.017) | 0.059 | 1.009 (1.001 to 1.017) | **0.033** |
| **Esophagus** | 1.001 (1.001 to 1.019) | **0.034** | 1.010 (1.002 to 1.018) | **0.013** |
| **Small Intestine** | 1.010 (1.000 to 1.021) | 0.054 | 1.011 (1.001 to 1.020) | **0.027** |
| **Liver** | 1.012 (0.998 to 1.026) | 0.089 | 1.013 (1.000 to 1.026) | 0.055 |
| **Stomach** | 1.000 (1.000 to 1.019) | **0.049** | 1.010 (1.001 to 1.019) | **0.028** |
| **Lung** | 1.010 (1.000 to 1.020) | **0.031** | 1.010 (1.002 to 1.019) | **0.016** |
| **Skin** | 1.009 (0.997 to 1.021) | 0.111 | 1.010 (0.999 to 1.021) | 0.068 |
| **Total** | 1.001 (1.000 to 1.003) | 0.058 | 1.001 (1.000 to 1.002) | **0.029** |

Abbreviations: HR = hazard ratio; CI = confidence interval

### Clinical association of tissue-specific alloreactivity potential

The simulated T cell counts are summarized in Supplementary Table 3, where for every organ (and overall) the mean simulated T cell counts were larger in patients who eventually developed any form of GVHD, than they were in patients who did not develop GVHD (P=N.S.). In unadjusted Cox proportional hazard models, esophagus-, stomach-, and lung-specific simulated T cell counts were significantly associated with cumulative GVHD incidence (P-values 0.034, 0.049, and 0.031 respectively; Table 2). When adjusted for recipient age and gender, the associations observed were also found to be significant for total and all organ-specific T cell simulation scores except for skin and liver (Table 2). This observation implies that within age and gender groups, the alloreactive T cell simulations may potentially identify donors with a higher likelihood of precipitating alloreactivity.

Separate unadjusted Cox proportional hazards models did not find any association between total or organ-specific T-cell counts and acute or chronic GVHD. After adjusting for age and sex the results did not change except for the association between small intestine and chronic GVHD (Supplementary Table 4).

## Discussion

Alloreactivity following SCT is a complex disorder dependent on numerous of factors such as degree of HLA matching, [28, 29, 30] intensity of immunosuppression [31, 32, 33] and the graft source/T cell composition of the graft [34, 35]. However, even within uniformly HLA matched and immunosuppressed patients with similar disease biology, outcomes remain variable and subject to laws of probability. In this paper the magnitude of difference in putative alloantigen burden and predicted T cell response in SCT DRP is explored, accounting for the HLA make up and protein coding differences between individual donors and recipient. Logically, the binding affinity and tissue expression of the antigens are critical parameters when assessing the T cell response to these antigens. The findings of an earlier model are now extended to simulate T cell responses to an array of alloantigens that the donor T cells may encounter in different organ systems in the recipient to determine tissue-specific alloreactivity-potential. Similar to the previous study, the findings reported herein demonstrate a very large degree of variability between the simulated T cell responses to antigens expressed in specific organs between unique DRP. Furthermore while, given the small sample size, and uniform simulation parameters applied to a heterogeneously treated cohort of patients, the association between GVHD in specific organs and the simulated T cell responses to the alloreactivity operator for these organs is not established, there are intriguing associations observed. Despite the use of anti-thymocyte globulin in a number of patients, cumulative GVHD was weakly associated with the calculated T cell responses when adjusting for recipient age and gender. While these are weak associations, it is remarkable that they were identified in this small cohort. This method of T cell response simulation represents a new potential area of investigation, which may yield interesting results in the future.

The inability to identify stronger clinical association between the magnitude of the simulated CD8+ T cell response to mHA array in a DRP and organ-specific GVHD, may be explained by, (a) uniform simulation conditions applied in the model, despite a heterogeneously treated patient cohort, with limited numbers in each group, (b) only 9-mer oligo peptides bound to HLA class I considered in the simulations, and those too without any post translational modification (phosphorylation, glycosylation) and protein cleavage site information incorporated and (c) our input data being derived from ‘normal’ tissue expression levels from GTEX, as opposed to patients potentially having different gene expression profiles in the post transplant state. Beyond these obvious considerations there are other important considerations that may be factored in mathematically in future iterations of this work, these are discussed below.

Logic would dictate that when proteins are cleaved, there will be a host of peptides generated for antigen presentation, both alloreactive, such as the ones reported here, as well as non-alloreactive peptides which will also bind HLA with varying levels of affinity, as the alloreactive peptides do. Furthermore these peptides will likely be far more numerous than the alloreactive ones, and may constitute a significant competitive barrier to the presentation of alloreactive peptides. This competition may be modeled numerically using the notion of *combinatorial probability* [36, 37]. In other words, if there are *n* total peptides which might be presented on *k* HLA molecule there are,

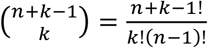
possible combinations of peptides which might be presented by these HLA molecules, including duplication of peptides and without regard to order of peptides presented. For example, if 10 peptides are to be presented by 4 HLA molecules the total number of possible peptide-HLA combinations will be, 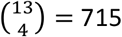. Now, of these, if 3 peptides are alloreactive (mHA), the probability of a favorable outcome (no GVHD) will be enhanced if the other 7 non-alloreactive peptides are presented, the number of possible combinations allowing that will be 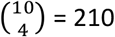. Therefore the probability of having a peptide combination presented by the four HLA molecules, which will not include an alloreactive peptide, i.e. a mHA in this mix of non-alloreactive and alloreactive peptides, will be approximately

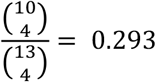
This implies that the remaining ^∼^70% of the peptide-HLA combinations will contain one or more mHA, if this combination of alloreactive and non-alloreactive peptide combinations were to be presented. Thus in the scenario presented above, depending on the HLA binding affinity of the peptides and its tissue expression, and the presence of a T cell clone recognizing that specific mHA, alloreactivity may not be triggered in nearly a third of the instances. In reality, however, the number of non-alloreactive peptides is likely high resulting in a smaller ratio of alloreactive peptides being presented. Supporting this line of reasoning is the observation that expression levels of HLA molecules have been shown to impact alloreactivity incidence [38, 39]. This quantitative consideration adds an important variable to determining the likelihood of alloreactivity developing not accounted for in the analysis presented in this paper; i.e., the variability in the HLA binding affinity of the many non-alloreactive peptides. Further variability in this system is introduced by the relative abundance of each protein from which the peptides are derived. This too will modify the likelihood of peptide binding to HLA, due to competition. This discussion demonstrates that while dynamical systems modeling may reproduce the T cell frequency distribution patterns seen in patients, the randomness of the exome mutations, and subsequent peptide-HLA interactions makes the development of alloreactivity a *stochastic*, dynamical process. Hence, while patterns of reactive immune responses may be mathematically described, each of these patterns is associated with a certain probability of occurrence and of causing the associated clinical outcome. In other words, HLA matched patients with a higher simulated T cell count will likely be at greater risk of GVHD under certain uniform conditions of immunosuppression and vice versa.

Another source of difference between an idealized simulation and the ‘real world’ will be the interaction between APC and T cells. Both cell populations proliferate in response to inflammatory stimuli, with the APC proliferation driving T cell growth. This interaction too may be modeled, using the matrix based vector operator model. The results reported above are based on the notion that the APC matrix operator (**M**_APO_) has a constant effect on the donor T cell vector, in other words antigen presentation is a constant function of time, however in reality this is not the case. APC themselves respond to inflammatory signals and as previously demonstrated, follow logistic growth after stem cell transplantation, consequently antigen presentation follows a crescendo-decrescendo course over time (particularly with infections) [40, 41]. This model may be built into this system of vector operator immune response modeling. In this instance consider the scenario; tissue injury following conditioning therapy ensues, following which there is uptake of recipient antigens by the transplanted APC (presumably of donor origin) proliferating in response to the cytokines released because of the insult. The proliferating APC take up antigen, present the mHA (and non-mHA) and migrate to the regional lymph nodes. As these monocyte populations arrive in the regional nodes, the T cells corresponding to the mHA start proliferating and migrate out of the lymph node and home in on the target tissue, where they now induce further inflammatory injury, further expanding the APC proliferation, until the APC population reaches a steady state. Once the tissue damage is complete (or infection resolved), the APC will decline and a steady state memory T cell complement left behind. This is a positive feedback loop which ‘self regulates’ as steady state values are reached for the APC population, with an eventual decline in the antigen presentation (as pathogen numbers decline, or maximal tissue injury runs its course). Similar to T cell clonal proliferation, APC proliferation may be described by equation 1. This allows the waxing and waning antigen presentation over time to also be described as a vector, and its impact in T cell population be recorded by taking the dot product of this APC effect vector and the T cell vector as it transforms. The APC effect vector, describing the impact of antigen presentation, as reflected by APC growth and decline on T cell clonal proliferation is given by the expression,

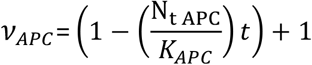

This variable is at its highest relative value, 2, in the beginning of the reaction (*N_t_* _*APC*_/*K*_*APC*_ approaches 0) but as the reaction proceeds over time *t*, it approaches 1, as *N*_*t APC*_ approaches *K* _*APC*_. Here 1 represents the homeostatic ability of T cell clone to persist in the absence of antigen presentation. Assuming that an independent *v_APC_* exists for each antigen in the M_APO_, the effect of APC growth and subsequent mHA presentation may be obtained by the dot product of the two vectors, *v_APC_* and *v_TCR_*. In other words, multiplying each iteration of the T cell vector with each iteration of this APC effect vector (the operation of multiplying 2 column matrices), transforms the donor T cell vector,

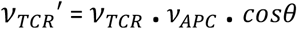

This equation implies that in the beginning of an inflammatory response there is a positive feedback response from proliferating APC, which amplifies the T cell response *v_TCR_ʹ*; eventually both decay to a steady state level. The angle *θ* between the two vectors is 0° (cos *0 = 1*) because they have the same direction (antigen specificity), in other words the TCR recognizes and binds to the mHA-HLA complex being presented by the APC. This effect of *v_APC_* on *v_TCR_* can be applied to the either a single T cell or the entire T cell repertoire (Figure 4A & B). As can be seen these graphs recapitulate the T cell response amplification commonly observed in response to antigen stimulation, [40, 41, 42] and can be depicted in the model illustrated in Figure 4C. In practice this interaction ‐‐ depending on the presence or absence of inflammation in the tissues being studied ‐‐ will significantly modify the alloreactivity operator and lead to variability in T cell proliferation observed. The weak associations of the T cell simulations with the clinical findings reported in this paper, may therefore be explained by absence of information regarding the overall inflammatory state in each DRP. This mechanism may also underlie the sensitivity of transplant outcomes to initial conditions and the seemingly chaotic outcomes of transplant dynamical systems [43].

**Figure 4.**
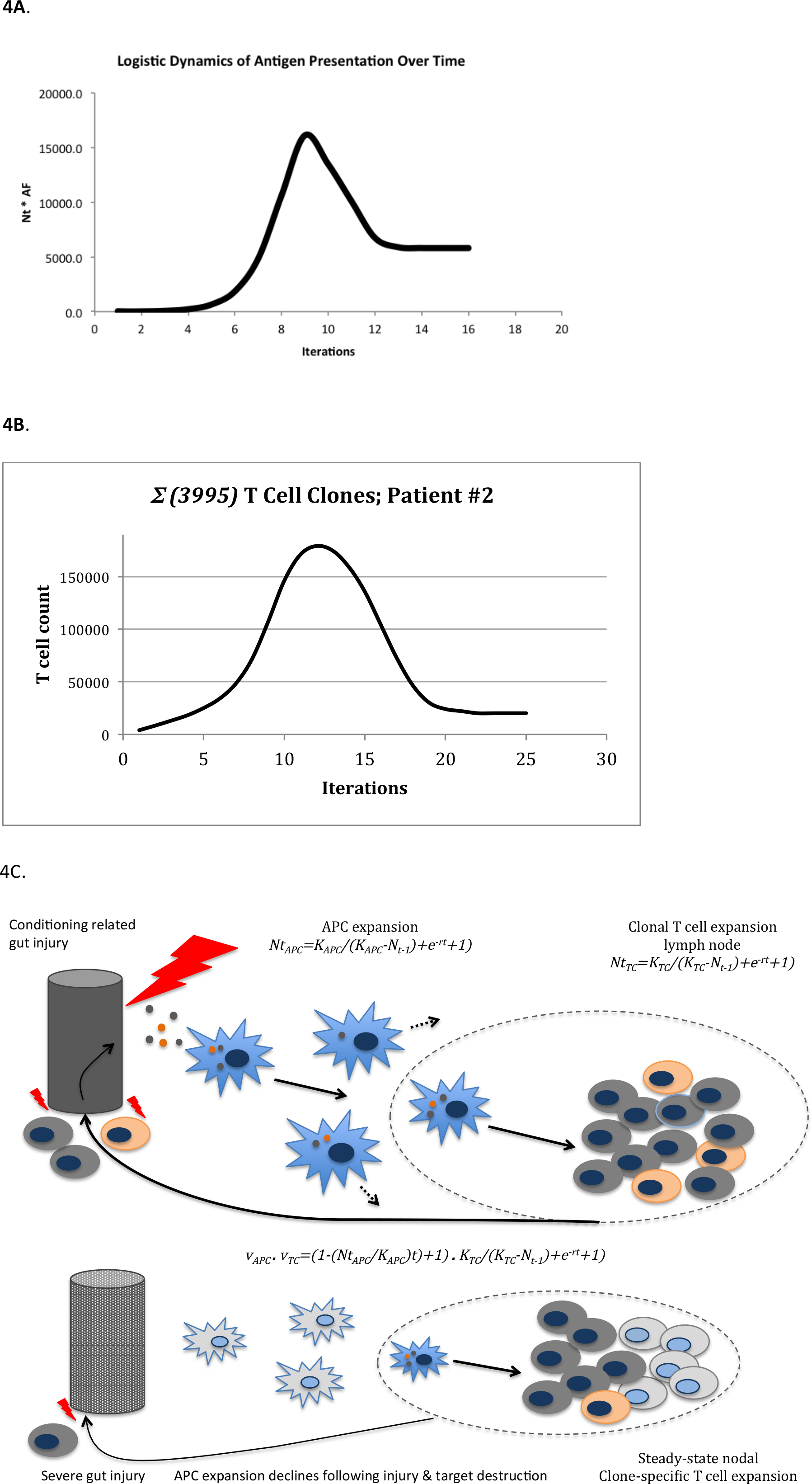
Alloreactivity model. Dot product of APC and T cell vectors recapitulates familiar antigen-challenge driven T cell proliferation response curve. A. Single T cell clone, B. Entire repertoire. C. Model illustrating the interaction between APC and T cells.

Another source of clinical variability not accounted for in this model is the constitution of the graft both in terms of effector and regulatory T cells (Treg). While the notion of absence of antigen directed clonal T cells impacting the predictive power of any such model is straightforward, the effect of regulatory immune cell populations such as Treg requires some calculation to understand. Treg recognize self-antigens and secrete anti-inflammatory cytokines (IL-4, IL-10) in response, diminishing the proliferation of the antigen directed CD8+ effector T cells [44]. This may be easily accounted for in this model by considering that the growth parameter *r* (given the uniform value of -1.5 in this analysis) represents the proliferative effect of the cytokine milieu. In this instance the more negative its value (induced by pro-inflammatory cytokines), the larger, and more rapid the growth promoting effect; and for less negative or alternatively for positive values (anti-inflammatory cytokine effect) growth will be impeded or stopped, as can be seen in Figure 5, where a change in the magnitude of *r* produces dramatic decline in the T cell growth curves. Once again these effects are not accounted for in the model presented here but may be easily included. Nevertheless, the robustness of the vector operator model of quantifying T cell response and its eventual clinical utility are indicated by its robustness in quantifying a variety of immune phenomenon noted above.

**Figure 5.**
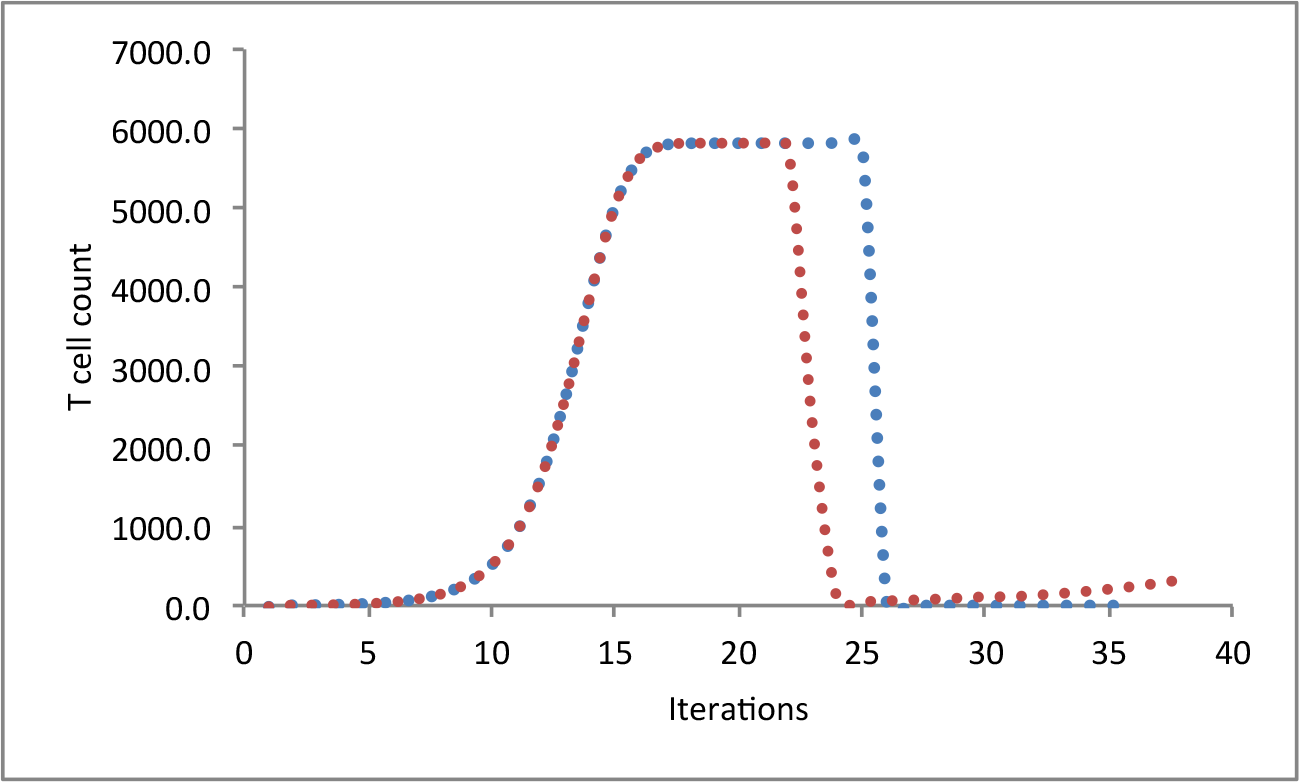
Modeling the effect of Treg on effector T cell growth: in the red curve, *r* reduced at 21st iteration from -1 to -0.25; T cell population drops but then recovers slowly. In the blue curve r reduced at 25th iteration from -1to +0.25 with direction reversal (from – to +), signifying anti-inflammatory cytokine effect supersedes pro-inflammatory cytokine effect.

Current clinical practice is to identify the donors based on the degree of major histocompatibility locus matching. Based on this, immunosuppressive regimens are chosen; unrelated donor transplant recipients frequently receive ATG in addition to the standard calcinuerin inhibitor + methotrexate, whereas haploidentical transplant recipients receive post-transplant cyclophosphamide. In the absence of early GVHD onset, immunosuppression intensity is gradually reduced and eventually withdrawn over a predetermined period of time, usually spanning four to six months, depending on the relapse risk of the underlying malignancy. Dynamical system modeling of immune reconstitution, based on whole exome sequencing data from donors and recipients may enable the development of patient specific immunosuppressive regimens post-transplant, by more precisely calibrating the GVHD risk that unique donors might pose to that patient. Our data indicate that adjusting for recipient biology this may be a powerful tool for identifying unique DRP that are at higher then average risk of GVHD and may need more measured adjustment of immunosuppression following SCT than others.

Given the advances in *next generation sequencing*, as well as computing technology it is conceivable that in time, patients and their prospective donors may have their HLA type and alloreactivity potential determined by whole exome sequencing. A donor with optimal alloreactivity potential will be identified and a GVHD prophylaxis regimen of optimal intensity utilized to achieve maximal *likelihood* of good clinical outcome. In so doing, dynamical systems understanding of alloimmune T cell responses will *attenuate* the unpredictability that the current largely probability-based models of outcomes prediction are fraught with. Dynamical systems analysis of antigenic variation also explains the randomness at hand in human immune response to disease, either infectious or neoplastic. This understanding has the potential to impact areas of investigation beyond transplant alloreactivity, potentially influencing cancer immunotherapy, autoimmune disease and infectious disease. Randomness within the dynamical system framework will remain always have to be accounted for because of the several mechanisms outlined above, but a model such as this is a step towards gaining a quantitative understanding of complex immune responses and their simulation to aid in transplant donor selection.

In conclusion this model partially explains why immune responses are seemingly random and difficult to accurately predict, akin to the quantum uncertainty principle, you can accurately measure either a particle’s position or its velocity, never both. Similarly, given the complexity at hand in immune responses, while we may not be able to precisely quantify the likelihood of alloreactivity, we hope that simulations of alloreactive T cell responses will aid the quantitative understanding of immune reconstitution and more closely estimate clinical outcomes.

---

**Supplementary Table 1.** Patient characteristics, excluding haploidentical DRP

|  |  |  |
| --- | --- | --- |
| **N** |  | 77 |
| **Male/Female** |  | 43/34 |
| **Age - median (range)** |  | 55.6 (21-73) |
| **Donor (%)** | MRD | 34 |
|  | MUD | 53 |
|  | MMRD | 1 |
|  | MMUD | 12 |
| **Stem Cell Source (%)** | Bone Marrow | 9 |
|  | Peripheral Blood | 91 |
| **Race**<br>**(Donor/Recipient)** | C/C | 50 |
|  | AA/AA | 13 |
|  | C/AA | 4 |
|  | MR/C | 2 |
|  | UK/C | 8 |
| **Conditioning Regimen (%)** | Reduced Intensity | 46 (60) |
|  | ATG/TBI | 25 |
|  | Busulfan/Fludarabine | 35 |
|  | Fludarabine/Melphalan | 3 |
|  | Myeloablative | 31 (40) |
|  | Busulfan/Cyclophosphamide | 22 |
|  | Cyclophosphamide/TBI | 14 |
|  | Etoposide/TBI | 1 |
| **GVHD prophylaxis (%)** | Cyclosporin A/Methotrexate | 21 |
|  | Cyclosporin A/MMF | 5 |
|  | Tacrolimus/Methotrexate | 44 |
|  | Tacrolimus/MMF | 30 |
|  | Anti-thymocyte globulin (ATG) | 81 |
| **Gender**<br>**Donor/Recipient** | M/F | 17 |
|  | F/M | 16 |
|  | M/M | 27 |
|  | F/F | 17 |

**Supplementary Table 2.** Simulated T cell counts per donor type by organ (presented in 1,000s of cells)

|  | MRD (N=27) | MUD (N=45) |
| --- | --- | --- |
|  | Mean (SD) | Mean (SD) |
| Salivary Glands | 13.8 (14.9) | 14.0 (20.0) |
| Colon | 18.4 (18.1) | 20.2 (30.1) |
| Esophagus | 17.2 (17.8) | 18.6 (28.1) |
| Small Intestines | 19.1 (20.3) | 19.9 (25.7) |
| Stomach | 19.6 (19.5) | 21.1 (27.3) |
| Liver | 18.4 (17.8) | 22.6 (32.1) |
| Lung | 12.9 (12.9) | 16.5 (25.6) |
| Skin | 19.3 (19.6) | 20.9 (25.9) |
| Total | 138.8 (139.3) | 154.8 (210.5) |

**Supplementary Table 3.**
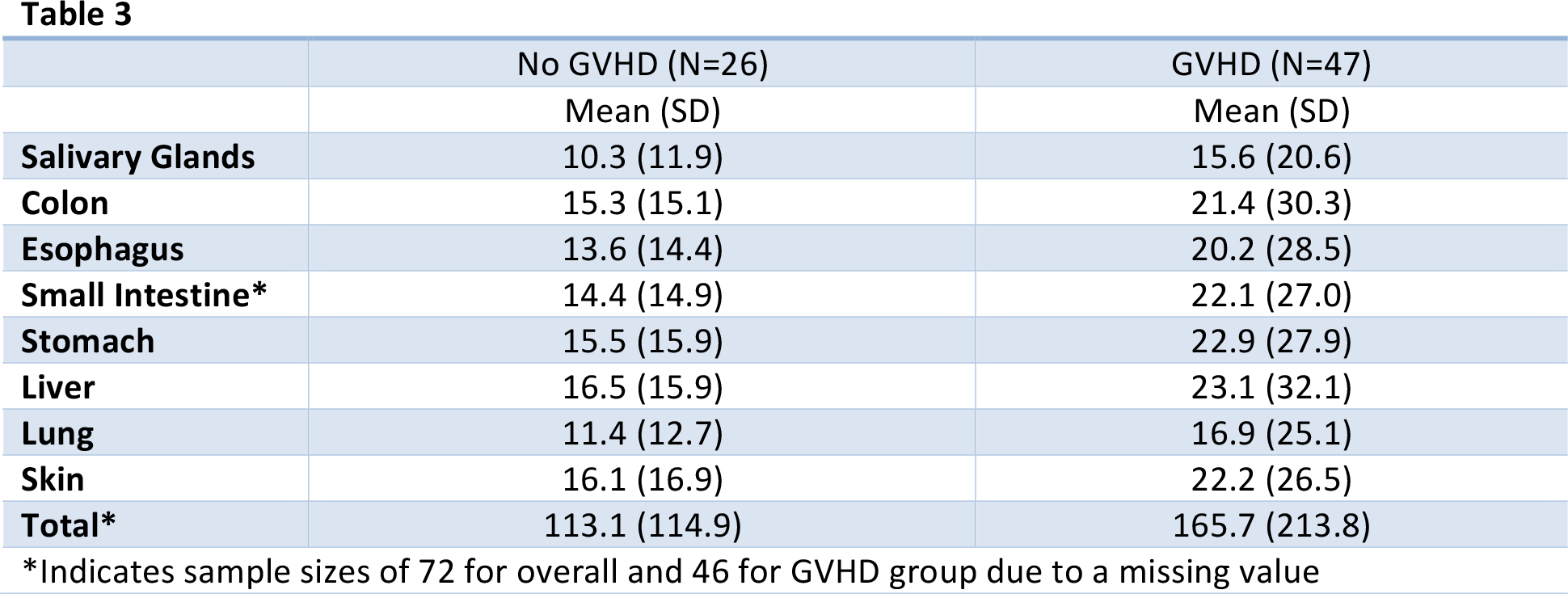
Simulated T cell counts by organ in patients with and without GVHD (presented in 1,000s of cells)

**Supplementary Table 4.** Estimated hazards ratios (HR), 95% confidence intervals (CI) and p-values from Cox-proportional hazards models of simulated T-cell counts and acute and chronic GVHD classification after adjustment for age and sex.

|  | Acute GVHD |  |  | Chronic GVHD |  |  |
| --- | --- | --- | --- | --- | --- | --- |
|  | HR* | 95% CI | p-value | HR* | 95% CI | p-value |
| **Salivary Glands** | 1.010 | 0.993, 1.027 | 0.250 | 1.014 | 0.998, 1.030 | 0.086 |
| **Colon** | 1.006 | 0.995, 1.018 | 0.274 | 1.007 | 0.995, 1.019 | 0.233 |
| **Esophagus** | 1.008 | 0.996, 1.019 | 0.177 | 1.009 | 0.997, 1.021 | 0.146 |
| **Small Intestines** | 1.007 | 0.993, 1.021 | 0.300 | 1.011 | 1.000, 1.023 | **0.049** |
| **Stomach** | 1.006 | 0.992, 1.020 | 0.391 | 1.010 | 1.00, 1.021 | 0.078 |
| **Liver** | 1.011 | 0.994, 1.028 | 0.206 | 1.014 | 0.997, 1.030 | 0.103 |
| **Lung** | 1.008 | 0.998, 1.019 | 0.103 | 1.009 | 0.99, 1.023 | 0.190 |
| **Skin** | 1.008 | 0.994, 1.021 | 0.278 | 1.009 | 0.99, 1.023 | 0.162 |
| **Total** | 1.001 | 0.999, 1.003 | 0.244 | 1.001 | 1.000, 1.003 | 0.119 |

**Supplementary Figure 1:**
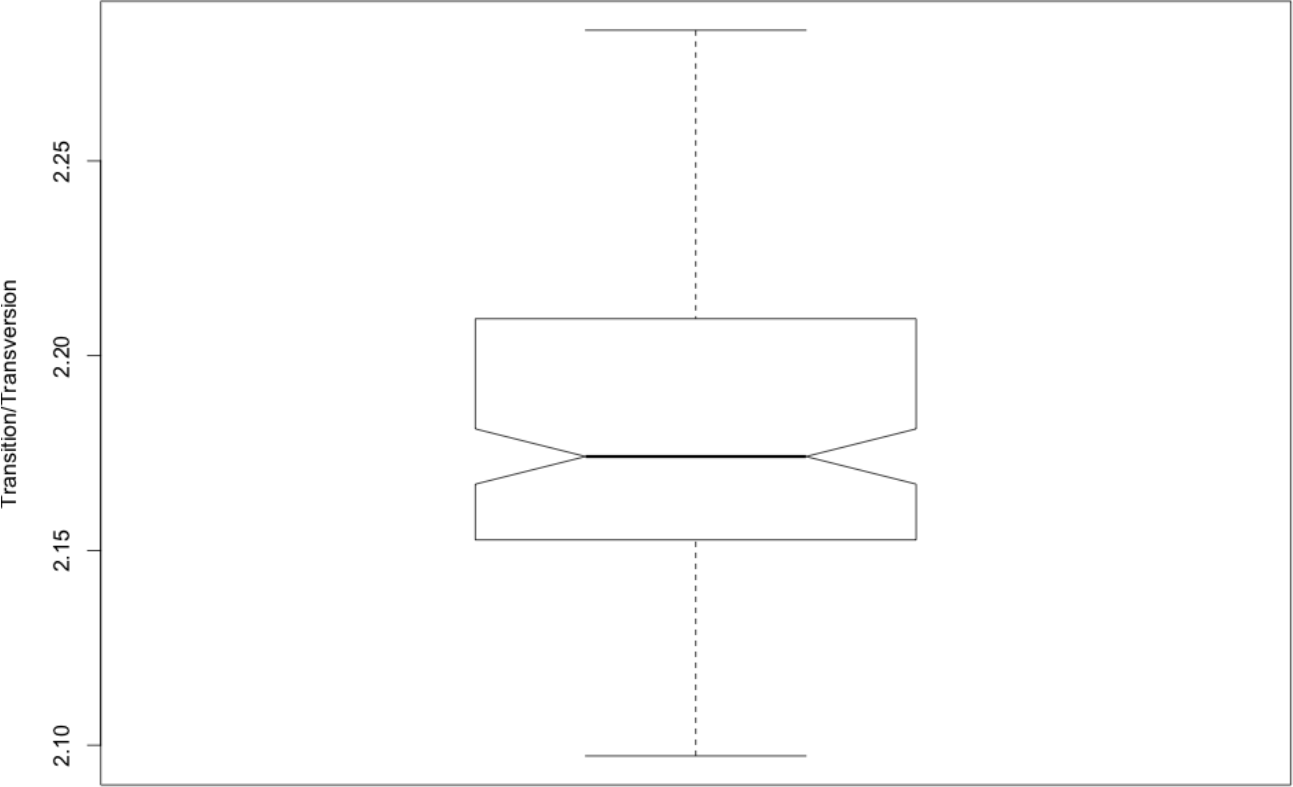
Transition/transversion ratio for the WES data from all the patients.

**Supplementary Figure 2:**
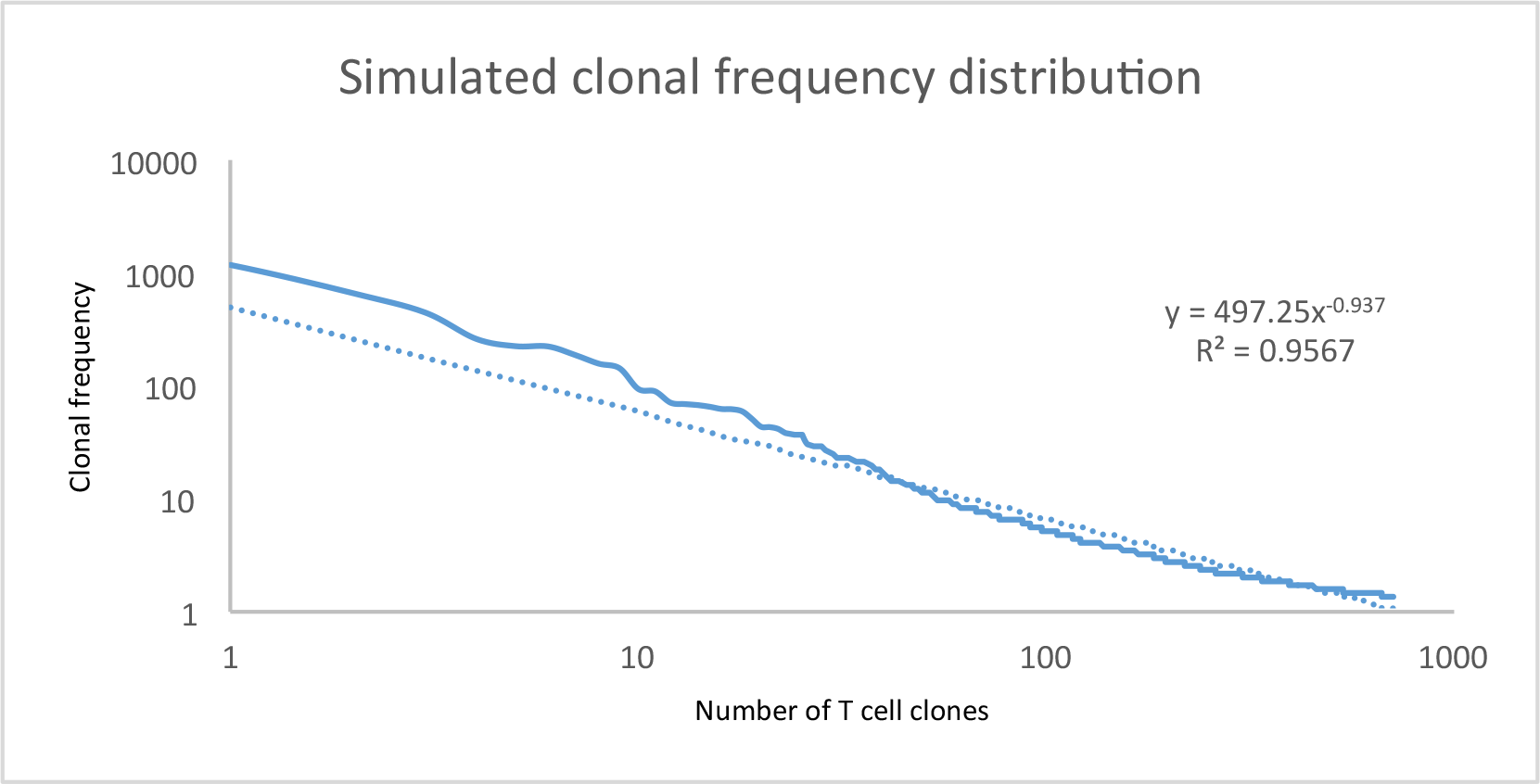
Clonal frequency distribution for colon responding T cells in patient 2.

## Supplementary Methods

If equation 3 is modified to balance the units, 
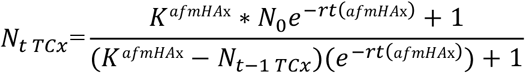
When solved this equation demonstrates near identical growth kinetics to those seen with Equation 3, however for the clones directed at low affinity, low expression *pmHA*, in other words, those clones with a low value of K, demonstrate erratic behavior not likely to be observed in reality. This suggests that in real life, thresholds of immune activation may be relevant and magnitude of T cell response important in immune reconstitution models such as this. For the calculations reported here the expression *e*^*‐rt*(*afmHA-x*)^+1 was not utilized in the numerator of the equation, and an *N*_*0*_ value of 1 was used.

## Acknowledgements

This work was supported by Massey Pilot Project Grant and an award from Virginia’s Commonwealth Health Research Board (PI: MN). VK performed bioinformatic analysis of the sequencing data to identify unique peptides and their HLA binding affinity, as well as tissue expression. BA wrote the program for performing calculations in MATLAB; BA and SS performed vector-operator calculations presented in this paper. AS, DJK and TS collected and verified clinical outcome data. RQ, AS and RS performed statistical analysis. MS performed sequencing on samples identified and procured by CR. AS, CH and MJ-L created data files with unique peptides and HLA IC50 values. All the authors contributed to writing the manuscript.

